# Morphological and biochemical responses of *Sphagnum* mosses to environmental changes

**DOI:** 10.1101/2020.10.29.360388

**Authors:** Anna Sytiuk, Regis Céréghino, Samuel Hamard, Frédéric Delarue, Ellen Dorrepaal, Martin Küttim, Mariusz Lamentowicz, Bertrand Pourrut, Bjorn JM Robroek, Eeva-Stiina Tuittila, Vincent E.J. Jassey

## Abstract

**Background and Aims:** *Sphagnum* mosses are vital for peatland carbon (C) sequestration, although vulnerable to environmental changes. For averting environmental stresses such as hydrological changes, *Sphagnum* mosses developed an array of morphological and anatomical peculiarities maximizing their water holding capacity. They also produce plethora of biochemicals that could prevent stresses-induced cell-damages but these chemicals remain poorly studied. We aimed to study how various anatomical, metabolites, and antioxidant enzymes vary according to *Sphagnum* taxonomy, phylogeny and environmental conditions.

**Methods:** We conducted our study in five *Sphagnum*-dominated peatlands distributed along a latitudinal gradient in Europe, representing a range of local environmental and climate conditions. We examined the direct and indirect effects of latitudinal changes in climate and vegetation species turnover on *Sphagnum* anatomical (cellular and morphological characteristics) and biochemical (spectroscopical identification of primary and specialized metabolites, pigments and enzymatic activities) traits.

**Key results:** We show that *Sphagnum* traits were not driven by phylogeny, suggesting that taxonomy and/or environmental conditions prevail on phylogeny in driving *Sphagnum* traits variability. We found that moisture conditions were important determinants of *Sphagnum* anatomical traits, especially those related to water holding capacity. However, the species with the highest water holding capacity also exhibited the highest antioxidant capacity, as showed by the high flavonoid and enzymatic activities in their tissues. Our study further highlighted the importance of vascular plants in driving *Sphagnum* biochemical traits. More particularly, we found that *Sphagnum* mosses raises the production of specific compounds such as tannins and polyphenols known to reduce vascular plant capacity when herbaceous cover increases.

**Conclusions:** Our findings show that *Sphagnum* anatomical and biochemical traits underpin *Sphagnum* niche differentiation through their role in specialization towards biotic stressors, such as plant competitors, and abiotic stressors, such as hydrological changes, which are important factors governing *Sphagnum* growth.

## Introduction

Northern peatlands are one of the largest stock of soil organic carbon (C) (Nichols and Peteet, 2019), and as such, have a major impact on the global climate cycle (Frolking and Roulet, 2007). Northern peatlands accumulated C over millennia due to cold, acidic, nutrients poor and water-saturated conditions that delayed litter microbial decomposition (Rydin and Jeglum, 2013). Bryophytes, and more particularly *Sphagnum* mosses, are vital for peatland C sequestration and storage as they effectively facilitate wet and acidic conditions that inhibits decomposition (Clymo and Hayward, 1982; Rydin and Jeglum, 2013; Turetsky, 2003). *Sphagnum* mosses also produce a recalcitrant litter and release antimicrobial biochemical compounds in the environment that further hamper the decomposition of dead organic matter (Fudyma et al., 2019; Turetsky, 2003; van Breemen, 1995). However, despite these general properties, we still lack a clear understanding of the species-specific characteristics, or traits, controlling *Sphagnum* growth and/or survival at global and local scale (Bengtsson et al., 2020a). *Sphagnum* species indeed show a high interspecific variability in their inhered anatomical and biochemical traits (Bengtsson et al., 2020b, 2018, 2016; Chiapusio et al., 2018; Dorrepaal et al., 2005; Gong et al., 2019). While *Sphagnum* anatomical traits are more and more used to explain how *Sphagnum* species adapt to environmental conditions (Bengtsson et al., 2020b; Jassey and Signarbieux, 2019), our knowledge on *Sphagnum* biochemical traits, how they vary among species and phylogeny, how they relate to anatomical traits and how they respond to local and global changes remain limited.

*Sphagnum* mosses produce thousands of metabolites, and can secrete hundreds of them in their surroundings (Fudyma et al., 2019; Hamard et al., 2019). Like in vascular plants, the role of these compounds in moss physiology and ecology can be either specific or multiple, but all of them play a role in moss fitness and/or tolerance to stress (Isah, 2019; Tissier et al., 2014). More particularly, a wide and important class of primary (carbohydrates, carotenoids) and specialized metabolites (phenols, proline, flavonols, tannins) have been suggested to play crucial roles in the growth, photosynthesis efficiency, litter-resistance to decomposition, and resistance to abiotic stresses of bryophytes (Xie and Lou, 2009). For instance, the production of carbohydrates and polyphenols can assist the maintenance of the integrity of plant cell-structure (i.e. anatomical tissues) and the internal regulation of plant cell physiology (Ferrer et al., 2008; Tissier et al., 2014). Furthermore, environmental transitory or constant environmental stress can cause an array of morpho-anatomical and physiological changes in *Sphagnum* mosses, which may affect *Sphagnum* growth and may lead to drastic changes in peatland functioning (Bengtsson et al., 2020b; Jassey and Signarbieux, 2019). In addition, environmental stress often leads to accelerated production of toxic metabolic by-products in plants, i.e. reactive oxygen species (ROS) (Chobot et al., 2008; Choudhury and Panda, 2005; Dazy et al., 2008; Liu et al., 2015; Roy et al., 1996; Sun et al., 2011; Zhang et al., 2017). ROS such as the superoxide anion (O_2_^•−^), hydrogen peroxide (H_2_O_2_), hydroxyl radical (OH^•^) can cause rapid damages to living tissues and macromolecules (e.g. DNA, proteins, lipids and carbohydrates) (Proctor, 1990), leading to an array of morpho-anatomical and physiological changes, which gain may lead to important changes in *Sphagnum* fitness and hence peatland functioning. In order to protect their cells against oxidative damages, mosses -like vascular plants- accumulate antioxidants such as flavonoids, tannins and/or produce antioxidant enzymes such as ascorbate peroxidase (APX), catalase (CAT), peroxidases (POX) and superoxide dismutase (SOD) (Choudhury et al., 2013; Das and Roychoudhury, 2014; Guan et al., 2016; Noctor et al., 2018). The regulation of the production of *Sphagnum* metabolites and of the activity of antioxidant enzymes are likely important processes in shaping *Sphagnum* morphology, growth, physiology and tolerance to environmental changes. Nowadays, limited number of studies focused on *Sphagnum* metabolites (Fudyma et al., 2019; Klavina, 2018), while antioxidant enzymatic activities remained unexplored. In addition, the linkages among *Sphagnum* morphological traits, metabolites and antioxidants are virtually unknown, while resource allocation among these different *Sphagnum* attributes could have important ramifications for peatland carbon cycling.

Here, we aimed to study how various anatomical metabolites and antioxidant enzymes vary according to *Sphagnum* taxonomy, phylogeny and environmental conditions. We conducted our study in five *Sphagnum*-dominated peatlands distributed along a latitudinal gradient in Europe, representing a range of local environmental and climate conditions. We examined the direct and indirect effects of latitudinal changes in climate, nutrient availability, and vegetation species turnover on *Sphagnum* anatomical and biochemical traits. We hypothesize that *Sphagnum* traits are strongly driven by phylogeny despite environmental changes. Previous works evidenced that production of metabolites in mosses can vary along microtopography, with vegetation changes (Binet et al., 2017; Chiapusio et al., 2018), depth (Jassey et al., 2011b), and environnemental conditions (Limpens et al., 2017). We, thus, expect that environmental changes to have a larger impact on *Sphagnum* traits than phylogeny. Furthermore, rates of antioxidant enzyme activities in bryophytes often vary with water deficit (Guan et al., 2016; Liu et al., 2015; Zhang et al., 2017) and low temperatures (Liu et al., 2017). Finally, we hypothesize that *Sphagnum* mosses will invest more in antioxidant metabolites and enzyme activities under drier climate than under wet conditions.

## Materials and methods

### Study sites and sampling

Five sites were selected along a latitudinal gradient spanning different environmental and climate conditions, from northern Sweden to southern France. Overall, these bogs covered a range of mean annual temperature from −0.1°C to 7.9°C (Figure 1a), and a range of precipitation from 419 mm to 1027 mm (Supplementary table S1). Counozouls is a poor fen in southwestern France that belongs to the Special Area of Conservation Natura 2000 site “Massif du Madres Coronat” (42°41’19.7“N 2°14’02.4”E) in the Pyrenees mountains. The moss layer was dominated by *Sphagnum warnstorfii* and *S. palustre* while vascular plants were dominated by *Mollinia sp*., *Carex sp*., *Ranunculus sp*. and *Potentilla anglica*. Kusowo Bagno is a bog located in northern Poland (53°48’47.9“N 16°35’12.1”E) in a nature reserve and is a part of the Special Area of Conservation Natura 2000 site “Lake Szczecineckie”. The peat moss layer was dominated by *S. magelanicum* and *S. fallax*, while *Eriophorum vaginatum*, *A. polifolia*, *V. oxycoccos*, and *D. rotundifolia* were the most common vascular plants at the sampling location. The Männikjärve bog (58°52’26.4“N 26°15’03.6”E) is located in Central Estonia in the Endla mire system. The ground layer was dominated by *Sphagnum capillifolium*, *S. fuscum* and *S. mefium* whereas *Drosera rotundifolia*, *Eriophorum vaginatum*, *Andromeda polifolia*, and *Vaccinium oxycoccos* characterized the vascular plant community. Siikaneva (61°50’41.6“N 24°17’17.5”E) is situated in an ombrotrophic bog complex in Ruovesi, Western Finland. The *Sphagnum* carpet was dominated by *Sphagnum fuscum* in hummocks and *S. papillosum*, *S. majus, and S. rubellum* in lawns-hollows. The vascular plant carpet was composed of dwarf shrubs (mainly *Vaccinium oxycoccos* and *Andromeda polifolia*), ombrotrophic sedges (*Eriophorum vaginatum* and *Carex limosa*) and herbs (*Scheuchzeria palustris* and *Drosera rotundifolia*). Abisko (sub-arctic Sweden; 68°20’43.1“N 19°03’58.7”E) is a gently sloping ombrotrophic peat bog surrounded by tundra. *S. balticum* was the numerically dominant peat moss species, whereas vascular plants such as *Carex sp*., *Andromeda polifolia, Rubus chamaemorus and Empetrum nigrum* were scarce.

**Figure 1.**
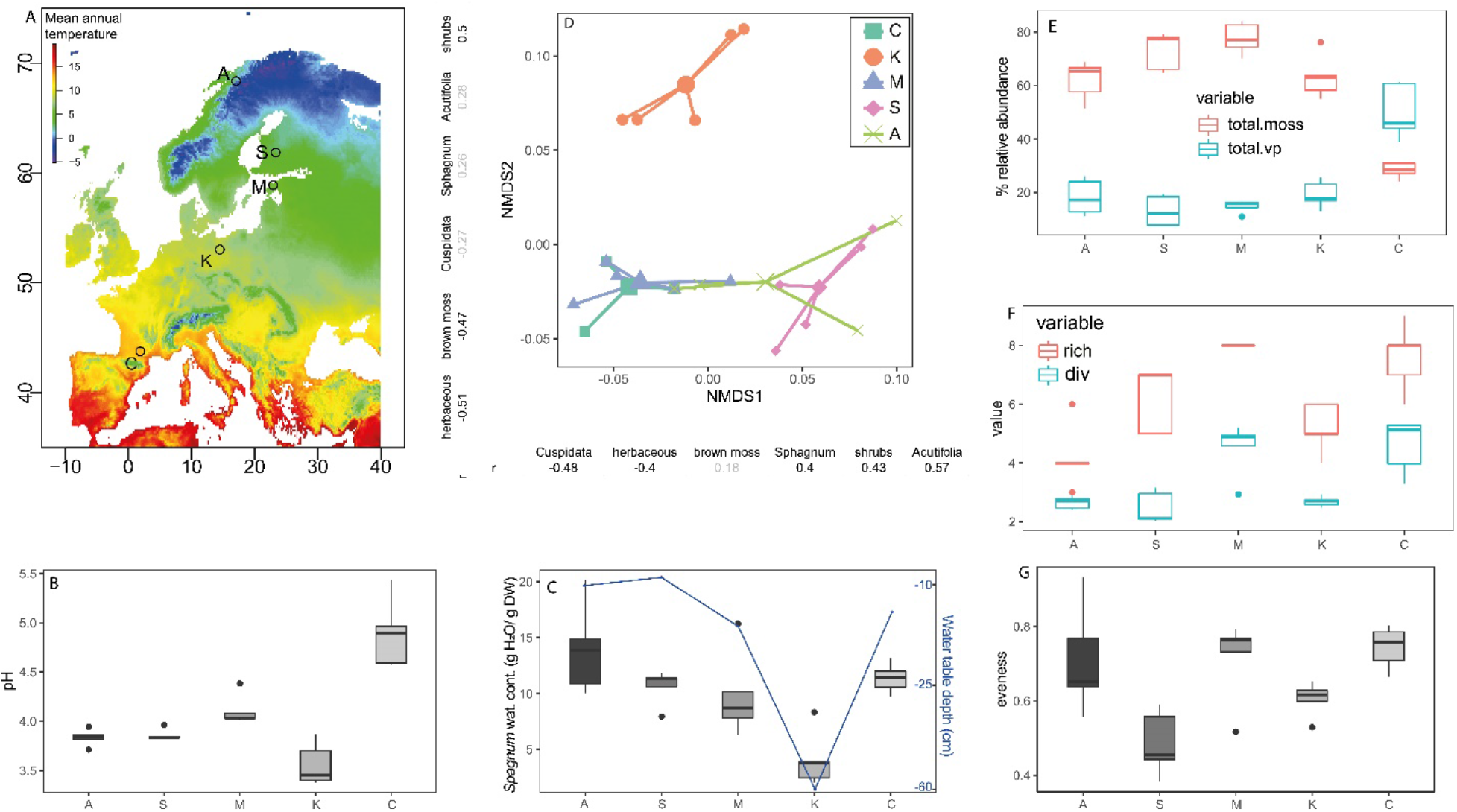
Locations and characteristics of latitudinal gradient. (a) Latitudinal gradient was established in five peatlands based on mean annual temperature. Boxplot (n = 5) showing mean (b) pore water pH, (c) *Sphagnum* water content (g H2O/g DW) and purple line represents water table depth (cm), (e) total vascular plant and *Sphagnum* abundance (%) of total moss and vascular plant (vp), (f) plant species richness (rich, S) and plant diversity (div., H’), (g) plant community evenness (J). (d) The two primary of non-metric multidimensional scaling (NMDS) based on Bray–Curtis distance showing differences in vascular plants and moss relative abundances in five sampling areas (stress=0.13). Sites are indicated with letters C=Counozouls, K=Kusowo, M=Mannikjarve, S=Siikaneva, A=Abisko

*Sphagnum* sample were collected at every site within the same week in early-July 2018. At each site, we selected the regionally dominant *Sphagnum* species based on a preliminary vegetation survey: *S.warnstorfi, S.magellanicum,, S. rubellum, S.papillosum and S. balticum.*(Supplementary table S2). Mosses were sampled at five homogeneous plots (50 × 30 cm each; 5 plots × 5 sites = 25 plots in total). Only the living *Sphagnum* tissue (top 3 cm from the capitula) were collected in plastic bag/ tubes according to the type of analyses, and kept in a cooler while travelling back to the laboratory. Samples were then either frozen at −20 °C or kept cold at 4°C depending on the type of analysis. Water-table depth (WTD) and groundwater pH were measured directly in the field using a ruler and a portable multimeter Elmetron CX742, respectively.

### Vegetation survey

We assessed the dynamic of plant species cover at each date in each plot using pictures taken in each plot (Supplementary table S2). Following Buttler *et al.* (2015), we took two high-resolution images in each plot (25 × 15 cm). On each picture, we had a grid of 336 points laid and quantified species overlaying the grid intersects. This technique did not support an estimation of vertical biomass distribution and possibly underestimated the frequency of certain species (Jassey et al., 2018) However, the potential bias was similar across plots, making species frequencies comparable among sites. We especially distinguished the dwarf-shrubs cover from the non-woody herbaceous cover as these two plant types differ in energy and nutrient allocation and litter production, and as such may have different effects on ecosystem C dynamic (Kruk and Podbielska, 2018).

### *Sphagnum* anatomical traits: cellular and morphological characteristics

We characterized the stress-avoiding adaptations to climate variation of each *Sphagnum* species using a suite of five anatomical traits, quantified following (Jassey and Signarbieux, 2019): volume of the capitulum (top layer), water-holding capacity of the capitulum and stem (0-1 cm), number of hyaline cells in tissues (i.e. dead cells storing water), and surface of hyaline cells (length x width), width of chlorocystes (bordering hyaline cells). In total, 25 individuals (5 per plot) were randomly collected to estimate the volume of the capitula (mm^3^) by measuring their height and diameter using a precision ruler. Then, we used the same samples to quantify the net water content at water saturated of the capitula and stem. Capitula and stems were submerged in water until their maximum water retention capacity was reached. Excess of water was removed by allowing water to flow naturally after removing capitula and stems from water. Then, individual capitula and stems were weighted as water-saturated and dried out after 3 days at 60°C. The net water content at water saturated of each individual was expressed in grams of water per gram of dry mass (g H_2_O_2_/g DW). At the cellular level, we randomly took 3 *Sphagnum* leaves from 125 capitula and prepared microscopic slides. We quantified the number of hyaline cells in tissues (number of hyaline cells per mm^2^), as well as their surface (μm^2^) using a light microscope connected to a camera (LEICA ICC50 HD). Three photos of 25 individuals from each five sites were taken (in total 375 pictures), and cell measurements made using the LEICA suite software.

### *Sphagnum* biochemical traits: primary and specialized metabolites, pigments and enzyme activities

Figure 2 shows the different extractions pathways used to quantify the various *Sphagnum* metabolites and enzyme activities. *Sphagnum* moss biomass was previously frozen, lyophilized and stored at −20°C before performing all biochemical analyses.

**Figure 2.**
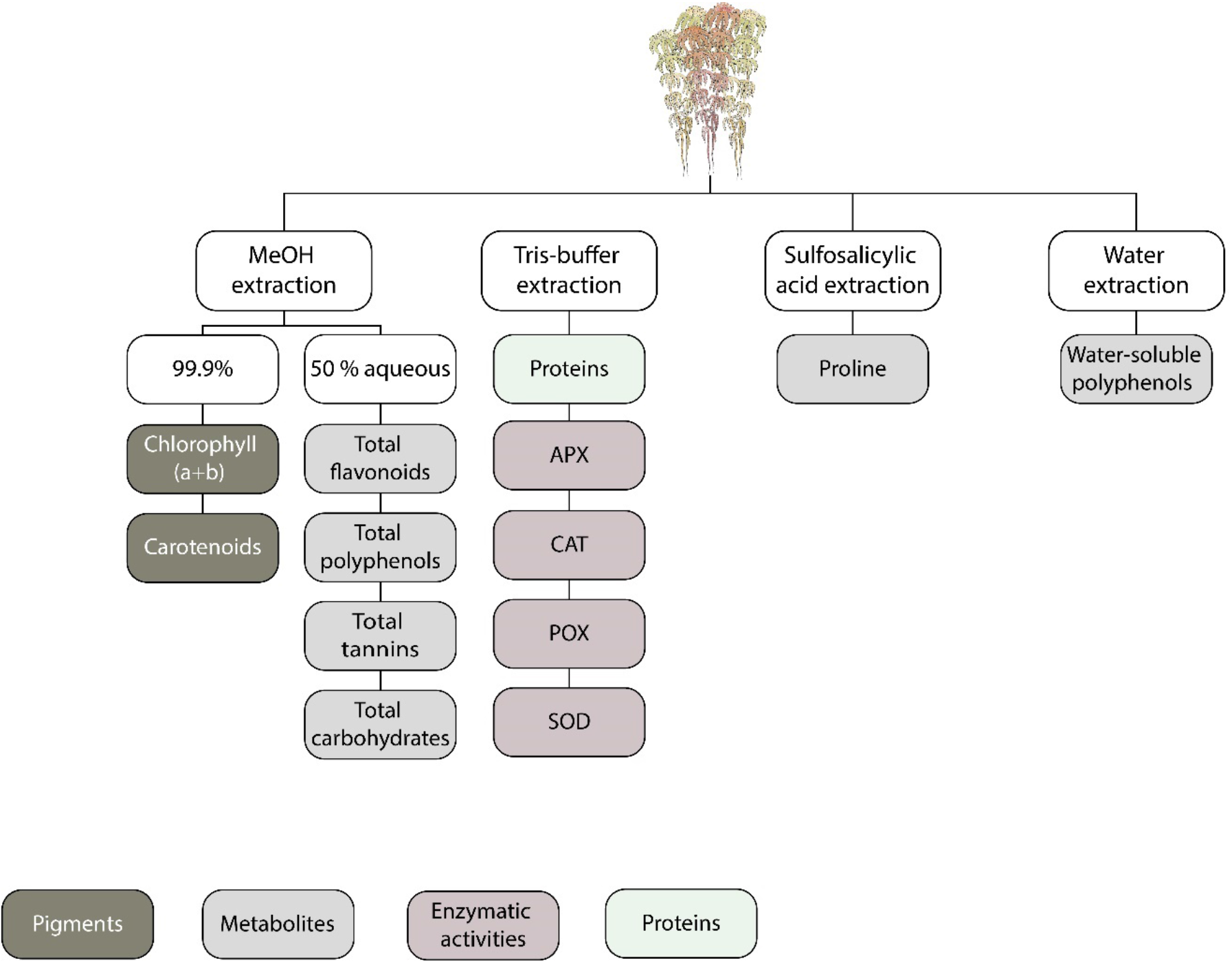
A summary of the extraction methods and compounds quantified in this study. Abbreviations are APX (EC 1.11.1.11) - ascorbate peroxidase activity, CAT (EC 1.11.1.6) – catalase activity, POX (EC 1.11.1.7) – peroxidase activity, SOD activity (EC 1.15.1.1) - superoxide dismutase.

### *Sphagnum* biochemical traits: pigments and metabolites

#### Pigments

Chlorophyll a, b and total carotenoids extractions were carried out in dark conditions. Twenty mg of lyophilized moss together with 1 mL of 99.9% cold methanol and 2 glass beads were placed into 2 mL tubes and then ground with high-speed benchtop homogenizer. A small spatula of MgCO_3_ was added to methanol in order to avoid acidification and the associated degradation of chlorophyll (Ritchie, 2006). Then, the tubes were centrifuged for 2 min at 8000 rpm at 4°C. The procedure of grinding and centrifugation was repeated two times in total. After that, the supernatant was transferred into 96-wells transparent microplate and the absorbance was measured with a spectrophotometer at 480, 652, and 665 nm. Concentrations of chlorophyll a, chlorophyll b and total carotenoids were calculated according to the extinction coefficients and equations reported by (Porra et al., 1989). Finally, pigments concentration was expressed as milligram of pigments per g dry weight (mg g^−1^ DW).

#### Methanol extraction of *Sphagnum* metabolites

We followed Klavina et al. (2018) (with some modifications) to extract *Sphagnum* metabolites, including total polyphenols, flavonoids, tannins and carbohydrates (Figure 2). Briefly, we ground moss samples in a high-speed benchtop homogenizer (FastPrep-24™, MP Biomedicals, China), and then added 40 mg of moss in a brown glass bottle with 4 mL of cold 50% aqueous methanol solution. After that, samples were exposed to an ultrasound bath for 40 min and bottles were shaken in an orbital shaker for 18 hours at 150 rpm, followed by another 40 min of ultrasound. Finally, moss extracts were replaced into 2 mL Eppendorf and centrifuged at 4500 rpm for 5 min at 4°C. Collected supernatants were then used for further analyses.

#### Total carbohydrates

Total carbohydrates quantification followed (Klavina et al., 2018) with slight modifications. 20 μL of moss methanolic extract was diluted with 200 μL of distilled water in 2 mL Eppendorf tube. Then 200 μL of 5% phenol solution and rapidly 1 mL of 98% H_2_SO_4_ were added. After 10 min of incubation at room temperature, test tubes were carefully shaken and then left for 20 min more. Absorbance was read on the spectrophotometer at 490 nm. A standard curve was prepared with glucose, thus values were expressed in mg equivalent glucose per gram of dry moss (mg g^−1^ DW).

#### Total polyphenols

The Folin-Ciocalteu method (Ainsworth and Gillespie, 2007) was used to determine total polyphenol content in *Sphagnum* tissues. *Sphagnum* methanolic extract was transferred into 96-wells transparent microplate with 40 μL of 10% Folin-Ciocalteu reagent. Then the microplate was covered with aluminum and shaken for 2 min using vortex, 140 μL of 700 mM NaHCO_3_ was added in each well, and the microplate was covered and incubated at 45°C for 20 min. Finally, the absorption of each well was measured using a spectrophotometer (Biotek Synergy MX) at 760 nm wavelength. Gallic acid was used as standard, so total polyphenolic content was expressed in mg equivalent Gallic acid per gram of dry moss (mg g^−1^ DW).

#### Total flavonoids

We used trichloroaluminum assay to assess the concentration of total flavonoids in *Sphagnum* tissues (Settharaksa et al., 2014). Twenty-five μL of moss methanolic extract was placed into 96-well microplate with 10 μL of 5% NaNO_2_. Then, the plate was covered with aluminum foil and incubated for 6 min at 20°C. Following incubation, 15 μL of 10% AlCl_3_ and 200 μL of 1M NaOH were added in each well, the plate was covered and shaken for 1 min. The absorption was measured on the spectrophotometer at 595 nm wavelength. Сatechin was used as standard and thus values expressed in mg equivalent catechin per gram of dry moss (mg g^−1^ DW).

#### Total tannins

We used acid vanillin assay to quantify total tannins concentrations in *Sphagnum* tissues (Bekir et al., 2013). Fifty μL of moss methanolic extract was placed into 96-well microplate and then 100 μL of freshly prepared 1% in 7M H_2_SO_4_ was added. After 15 min of incubation in the dark at room temperature, the absorbance at 500 nm was measured. Сathehine was used a standard and the results were expressed in mg equivalent catechin per gram of dray moss DW (mg g^−1^ DW).

#### Total water-extractable polyphenolic compounds

The quantification of total water-extractable polyphenols followed (Jassey et al., 2011b) with some modifications. Twenty mg of lyophilized moss together with 5 mL of distilled water were placed into brown glass bottles and exposed to ultrasound in an ultrasound bath for 40 min. After that, test bottles were placed in a shaker for 3 hours at 150 rpm. Once shaking finished, tubes were centrifuged at 4500 rpm for 5 min at 4°C. Then, 1.75 mL of collected supernatant was transferred in brown glass bottle and 0.25 mL of Folin & Ciocalteu’s reagent was added. Test bottles were vigorously mixed with vortex and then 0.5 ml of 20% Na_2_CO_3_ was added and the vortex was repeated. Finally, we put samples in bain-marie at 40°C. After 20 min of incubation the absorbance on the spectrophotometer at 760 nm was measured. Gallic acid was used a standard, so total polyphenolic contents were expressed in mg equivalent gallic acid per gram of dry moss (mg g^−1^ DW).

#### Proline analysis

We used acid ninhydrin assay to determine total proline (Lee et al., 2018). Briefly, 25 mg of lyophilized moss was ground with high-speed benchtop homogenizer with 1 mL of 3% (w/v) sulfosalicylic acid and centrifuged at 4500 rpm for 5 min at 4°C. Then, 1 mL of collected supernatant was mixed with 2 mL of acid ninhydrin reagent (1.25 g of ninhydrin and 100 mL of acetic acid glacial) and glass tubes were boiled at 100°C. After an hour of boiling, the reaction was stopped by placing tubes on ice. Absorbance was read at 510 nm.). A standard curve was prepared with L-proline, thus proline content was expressed as mg of proline per g dry weight (mg g^−1^ DW).

### *Sphagnum* biochemical traits: antioxidant enzymes and proteins

We quantified the activity of four antioxidant enzymesduring abiotic stress conditions: ascorbate peroxidases (APX), catalases (CAT), total peroxidases (POX) and superoxide dismutases (SOD). To evaluate enzyme activities, 50 mg of grinded lyophilized moss was shaken in an automatic bead shaker with 1 mL of extraction buffer containing 100 mM Tris buffer pH 7.0, 0.5M EDTA, PVP 6 g/L, 0.5M ascorbate, β-Mercaptoethanol, industrial protease inhibitor cocktail (Sigma). Then the moss extract was centrifuged at 4500 rpm for 5 min at 4°C (Pourrut, 2008). Finally, the supernatant (hereafter ‘enzyme extract’) was collected and stored at −80°C for APX, CAT, POX, SOD enzyme activities measurements. Dosage of proteins was further determined according (Bradford, 1976), using bovine serum albumin (BSA) as a standard. Briefly, 10 μL of enzymatic extract and 190 μL of Bradford reagent were added to 96 wells microplate. Then microplate covered with aluminum foil was shaken in a shaker for 10 min at 150 rpm and finally, the absorbance was read at 595 nm. APX, CAT, POX and SOD were expressed as μmol per min per mg of proteins.

#### APX activity

was determined by adding 10 μL of enzyme extract to 230 μL of a reaction medium consisting of 0.5 M potassium phosphate buffer (Na_2_HPO_4_ + KH_2_PO_4_), pH 7, in a 96-wells UV-microplate. Then 15 μL of 10 mM ascorbate was added and microplates were covered with aluminum foil. At the last moment, 10 μL of 100 mM H_2_O_2_ was added, and the decrease of absorbance due to ascorbate oxidation was quickly measured during the first 5 minutes of reaction at 25°C. Enzyme activity was estimated using the molar extinction coefficients 290 nm, ε = 2.6 mM^−1^ cm^−1^ (Nakano and Asada, 1981).

#### POX activity

was measured by mixing 40 μL of enzyme extract and 180 μL of a reaction medium containing 31.25 mM potassium phosphate buffer, pH 6.8, and 10 μL of 250 mM pyrogallol in 96-wells microplate. At the last moment, 10 μL of 500 mM H_2_O_2_ was added, and the formation of purpurogallin was followed at 420 nm during the first 5 minutes of reaction at 25°C. Enzyme activity was estimated using the molar extinction coefficients of purpurogallin at 420 nm, ε = 2.47 mM^−1^ cm^−1^ (Maehly and Chance, 1954).

#### CAT activity

was determined by adding 10 μL of enzyme extract to 200 μL of a reaction medium consisting of 10 mM phosphate buffer, pH 7.8, in 96-wells UV-microplate. At the last moment, 10 μL of 100 mM H_2_O_2_ was added and the consumption of H_2_O_2_ was measured at 240 nm during the first 5 minutes of reaction at 25°C. Enzyme activity was estimated using the molar extinction coefficients 240 nm, ε = 39 M^−1^ cm^−1^ (Aebi, 1984).

#### SOD activity

was measured in shadowed conditions by adding 20 μL of enzyme extract to 190 μL of a reaction medium containing 50 mM potassium phosphate buffer, pH 7.8, 20 μL of 250 mM methionine, 1.22 mM nitrobluetetrazolium (NBT), and 10 μL of 50 μM of riboflavin in 96-wells microplate covered with aluminum foil. A first lecture of absorbance was performed on the spectrophotometer at 560 nm and then microplate was exposed to light at 15-W fluorescent lamp at 20°C in order to launch the reaction of blue formazana formation ((Pourrut, 2008)). After 10 min of incubation the microplate was covered with aluminum foil to stop the reaction, and a second lecture of absorbance was performed at 560 nm. One unit of SOD activity was defined as the amount of enzyme required to result in a 50% inhibition of the rate of NBT reduction, and thus SOD activity is expressed as unit per mg of protein.

### Statistical analyses

For each plot, we calculated plant richness (S) as the total number of species present, plant diversity as the Shannon diversity index (H’) and plant community evenness as Pielou’s index (J). Due to the non-normal distributions of the data (*i.e.* environmental conditions and vegetation data), non-parametric Friedman tests were performed to assess the effect of a latitudinal gradient on plant communities. The Friedman-Nemenyi multiple comparison test was applied for post-hoc analyses to bring out groups of data that differ among sites.

Analyses of variance (ANOVAs) were used to assess the effect of species (fixed effect) on *Sphagnum* attributes, and the effect of site on vegetation and environmental conditions. Tukey’s multiple comparison test was used for post hoc analyses of differences among the levels of the fixed effects in the final model. Assumptions of normality and homogeneity of the residuals were checked visually using diagnostic plots and a Shapiro test. Log or square root transformations were used when necessary in order to meet these assumptions.

We tested the presence of a phylogenic signal within anatomical and biochemical *Sphagnum* trait matrices by rejecting the null hypothesis that trait values for two species are distributed independently from their phylogenetic distance in the tree (Blomberg and Garland, 2002). Firstly, we built the phylogenic tree of our five *Sphagnum* species using the loci transfer RNA^Gly^ (trnG) from GenBank (Shaw et al., 2020, 2019, 2010; Weston et al., 2015). The FASTA sequences of the trnG gene were extracted from NCBI data base, and then assembled into a phylogenetic tree using the *phylo4d* function from the *phylosignal* R package (Keck et al., 2016). We tested the presence of a phylogenetic signal within traits using autocorrelation (Abouheif’s *Cmean* index) and on evolutionary mode (Blomberg’s K) principles. With each index, we could test a null hypothesis of a random distribution of trait values in the phylogeny (i.e. no phylogenetic signal).

We performed nonmetric multidimensional scaling (NMDS) to investigate patterns of variation in vegetation cover, anatomical and biochemical traits in five sites along the latitudinal gradient. For all multivariate analyses, a standardization was applied on every matrix to downweight the influence of dominant taxa, anatomical and biochemical traits (Legendre and Legendre, 2012). Pearson’s correlations were tested among the two first NMDS axes and main plant functional types to identify the dominant vegetation type along our latitudinal gradient. Only significant correlations with *r*>0.40 and *r*<−0.40 were considered as relevant.

Multiple factor analysis (MFA) was used to symmetrically link 1) the three Sphagnum attribute data sets, *i.e.* anatomical traits, metabolites and pigmetns and proteins and enzymatic antioxidant activities and 2) the tree *Sphagnum* traits matrices and the two matrices describing the enviro-climatological conditions (WTD, pH, *Sphagnum* water content, annual temperature and the highest temperature of the warmest quarter) and vascular plant cover (shrubs and herbaceous relative abundances) data. Shrubs and herbaceous cover were selected as they were identified as the main vascular plant functional types that significantly influenced vegetation composition along the gradient (see Figure 1 and results for more details). MFA was chosen because it permits the simultaneous coupling of several groups or subsets of variables defined on the same objects and to assess the general structure of the data (Escofier and Pagès, 1994). MFA was conducted in two steps. First of all, a PCA was done on each data subset, and then it was normalized by division of all its elements by the eigenvalue acquired from its PCA. After that, those normalized subsets were clustered to get a unique matrix and a second PCA was done on this matrix. RV-coefficient (Pearson correlation coefficient) was used to assess the similarity between data matrices (Josse et al., 2008). To carry out cluster analyses in MFA (according to Ward method), Euclidean distances of global PCA were used and the final dendogram was projected in the MFA ordination space. MFA allowed to identify the main differences in the data structure described by all variables. This helps in revealing the main dissimilarities among the groups and/or sites (Borcard et al., 2011).

Redundancy analysis (RDA) was used to assess the ordination patterns of the *Sphagnum* anatomical and biochemical traits (standardized transformation) and their causal relationship to environmental and vegetation data sets (ter Braak and Smilauer, 1998). Adjusted R^2^ was used to estimate the proportion of explained variance (Peres-Neto et al., 2006). The significance of each explanatory variable included in RDA was tested using 1,000 permutations. The analysis was performed with *Sphagnum* traits and environmental and vegetation variables data sets in order to check the effect the effect of environmental and vegetation variables on *Sphagnum* anatomical and biochemical traits.

## Results

### Environmental conditions and vegetation across sites

Overall, the five sites differed in terms of water chemistry. Water table depth (WTD) was the lowest in Kusowo (WTD = −60 cm) and the highest in Siikaneva (WTD = – 8 cm; Friedman test, χ2=20, *P* < 0.01; Figure 1c). *Sphagnum* water content was the lowest in Kusowo with 4.1 gH2O/g DW and the highest in Abisko with 13.96 gH2O/g DW (Friedman test, χ^2^=12, *P*=0.017; Figure 1b). The lowest pH was measured in Kusowo and the highest in Counozouls (Friedman test, χ^2^=17.12, *P* < 0.01; Figure 1b).

The overall vascular plant and *Sphagnum* species richness differed significantly among sites (Friedman test; χ^2^=18.55, *P*<0.001; Figure 1f). Abisko had the lowest richness with 4 species on average, and Männikjärve had the highest richness with 8 species. Plant diversity and eveness were the lowest in Siikaneva (2.5 and 0.49 respectivelly) and the highest in Counozouls (4.6 and 0.74 respectivelly; diversity: Friedman χ^2^=23.3, *P*=0.01; eveness: Friedman χ^2^=10.72, *P*=0.03; Figure 1f-g). Both *Sphagnum* and vascular plant relative abundances varied along the gradient (*Sphagnum*: *F*_4,16_=25.9, *P* < 0.001, ANOVA; vascular plants: *F*_4,16_=31.8, *P* < 0.001, ANOVA; Figure 1d). *Sphagnum* relative abundance was the lowest in Counozouls (~32%) and the highest in Männikjärve (~78%) and Siikaneva (~73%), while vascular plants showed opposite patterns. Futhermore, the plant community composition differed markedly across sites (Figure 1e). While no particular variation could be seen on the first NMDS axis, Kusowo was well separetd from the other four sites along the second NMDS axis. More specifically, NMDS axis 1 was characterized by a decreasing abundance of herbaceous species (*r* = −0.64) and *Sphagnum* from *Cuspidata* subgenus (*r* = −0.48), while *Sphagnum* species from *Sphagnum* subgenera (r=0.4), shrubs (r=0.43) and *Sphagnum* species from the *Acutifolia* subgenera (*r* = 0.57) all increased. NMDS2 axis revealed the decreasing abundance of herbaceous (r=−0.51), brown moss (r=−0.47, while srubs (r=0.5) increased (Figure 1e).

### Phylogenic and taxonomic effects on *Sphagnum* attributes

While *Sphagnum* attributes markedly differed among species (Figure 3 and 4), we did not evidence any particular phylogenetic signal in *Sphagnum* anatomical and biochemical traits (*P* >0.05 in most cases), neither in proteins and enzymatic activities (*P* > 0.05 in all cases; Supplementary table S3). Only the capitulum volume (*Cmean* = 0.16, *P* <0.05; Supplementary data S3) was significantly related to *Sphagnum* phylogenetic distances.

**Figure 3.**
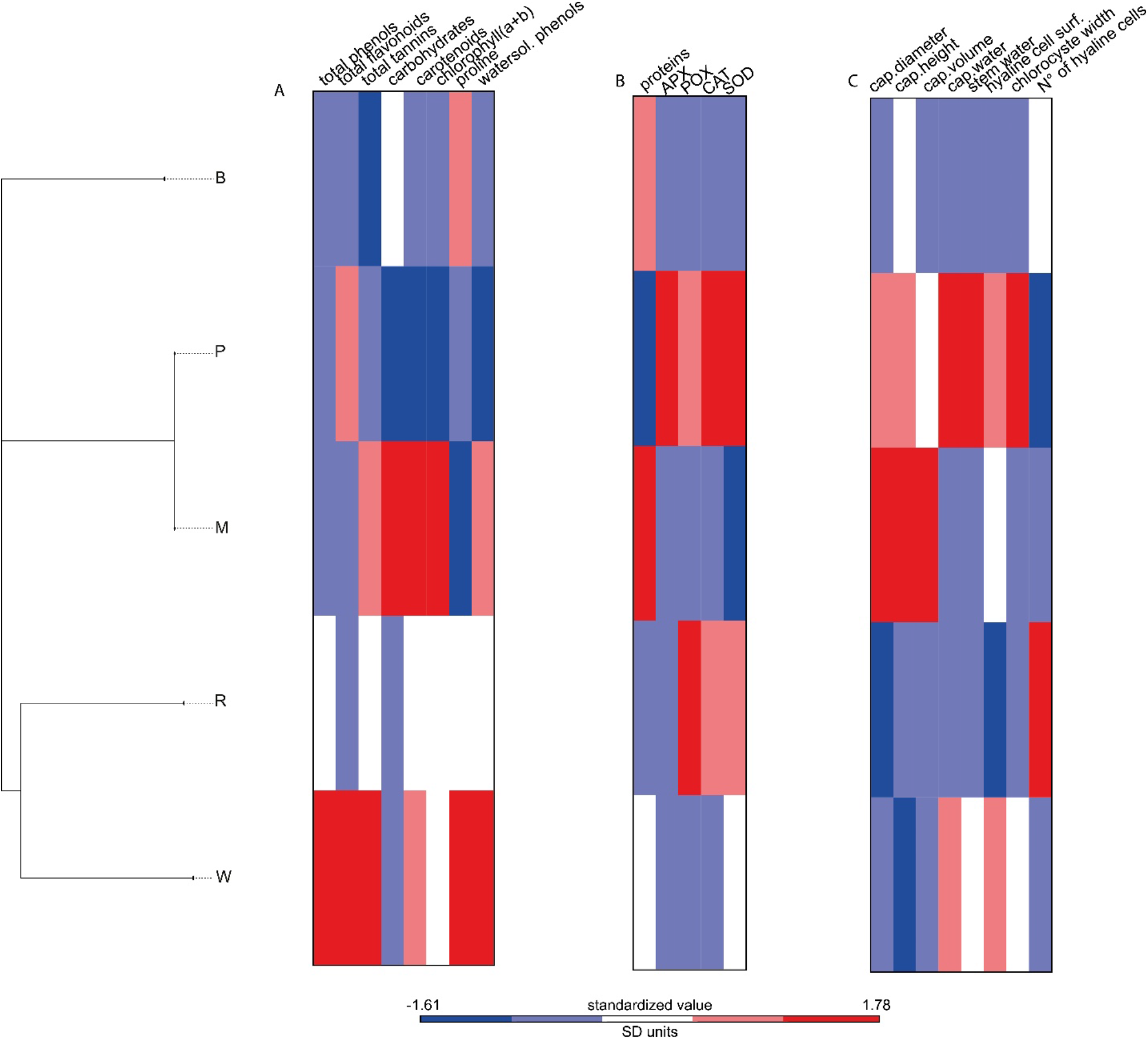
*Sphagnum* phylogenetic tree with the corresponding heatmap showing the dissimilarities in *Sphagnum* (A) metabolites and pigments concentrations (B) proteins and enzymatic activities, (C) anatomical traits. Colors represent the standardized value calculated from standardized means of *Sphagnum* attributes. *Sphagnum* species are indicated by letter W=*S. warnstorfii*, M=*S. magellanicum*, R=*S. rubellum*, P=*S. papillosum*, B=*S. balticum* followed by number indicating the plot number.

**Figure 2.**
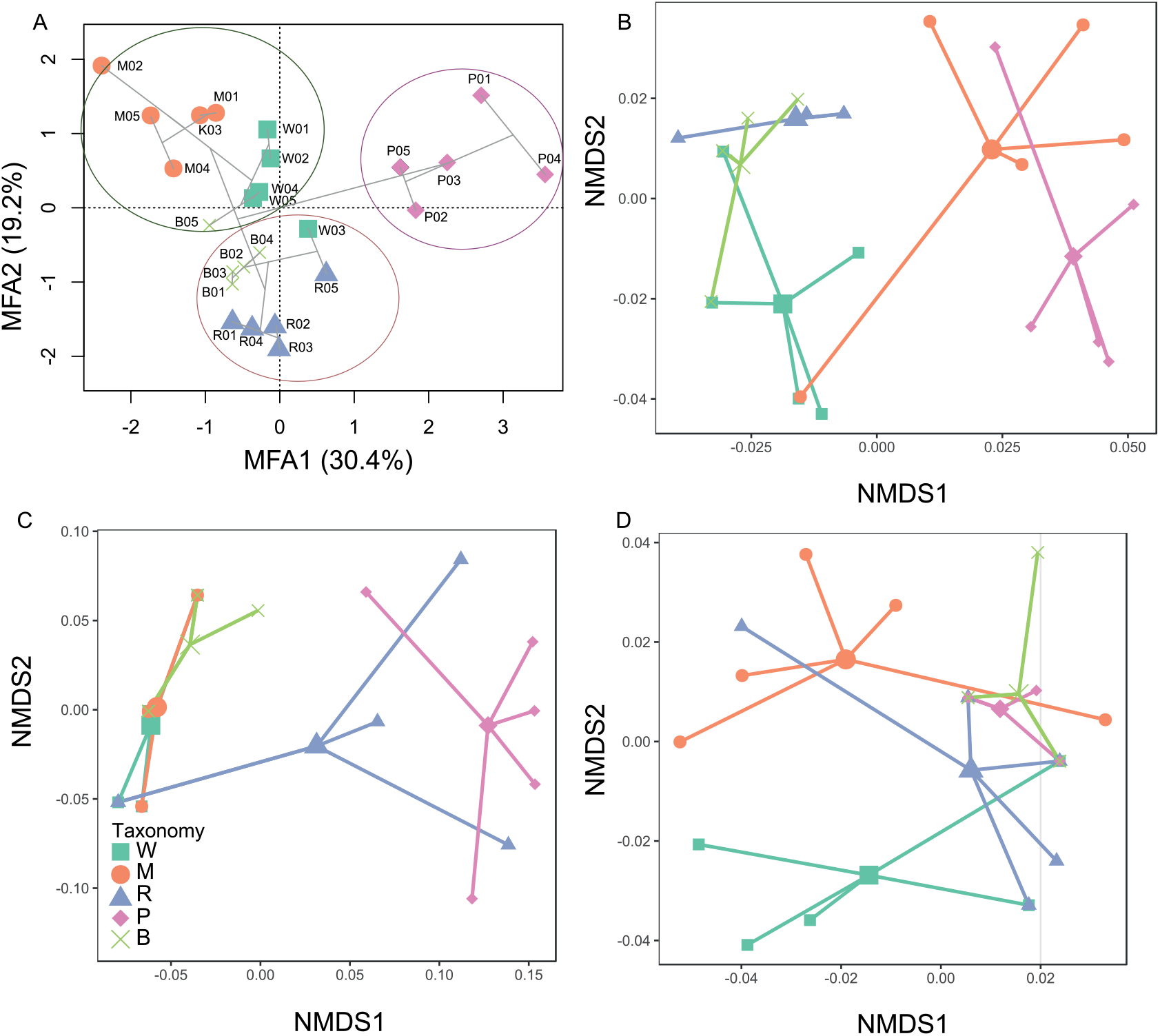
(a) Multiple factor analysis (MFA) of *Sphagnum* attributes (standardized data) data sets from five dominant *Sphagnum* species. Projection of the MFA axes 1 and 2 with the result of a hierarchical agglomerative clustering (grey lines), obtained by the Ward method on the Euclidean distance matrix between MFA site scores, showing four main groups. *Sphagnum* species are indicated by letter W = *S. warnstorfii*, M = *S. magellanicum*, R = *S. rubellum*, P = *S. papillosum*, B = *S. balticum* followed by number indicating the plot number. (b-d) The two primary of non-metric multidimensional scaling (NMDS) based on Bray–Curtis distance showing differences in *Sphagnum* (b) anatomical traits (stress=0.12), (c) proteins and enzymes (stress=0.13), and (d) metabolites and pigments (stress=0.177). Samples are represented by dominated *Sphagnum* species (different icons and colors).

When all *Sphagnum* attributes (i.e. the three data sets: metabolites and pigments, protein and enzymatic activities, and anatomical traits) are merged together, three distinct groups– not related to phylogeny– emerged: one comprised of *S. papillosum*, second of *S.balticum* and *S. rubellum*, third of *S. magelanicum* and *S. warnstorfii* (Figure 4a). The separate NMDSs on each specific trait category showed contrasting patterns (Figure 4b-d). For *Sphagnum* anatomical traits, two separate groups emerged along the first axis (Figure 4b): one group composed of *S. rubellum, S. warnstorwii* and *S.balticum* and a second group composed of *S. magellanicum* and *S. papillosum*. These patterns are explained by significant differences among individual anatomical traits (Figure 5). *S. magellanicum* had the highest capitulum diameter, capitulum height and capitulum volume with 1 cm^2^, 0.424 cm and 0.337 cm^3^ respectively, while the lowest values were observed in *S. rubellum* with 0.524 cm^2^, 0.25 cm and 0.006 cm^3^, respectively (diameter: *F*_4,16_=27.8, *P* < 0.001; heigh: *F*_4,16_=8.53, *P* < 0.01; volume: *F*_4,16_=23.15, *P* < 0.001, ANOVAs). *S. rubellum* had the highest number of hyaline cells (on average 693.2 mm^−1^) (*F*_*4,16*_=21.53, *P* < 0.001, ANOVA), but the highest water holding capacity was found in *S. warnstorwii* and *S.papillosum* (Figure 5; *F*_4,16_=10.61, P < 0.001, ANOVA). The width of chlorocystes was the highest in *S. papillosum* with 9.86 μm, while the other species had about two-fold lower width (Figure 5; *F*_4,16_=26.41, *P* < 0.001, ANOVA).

**Figure 3.**
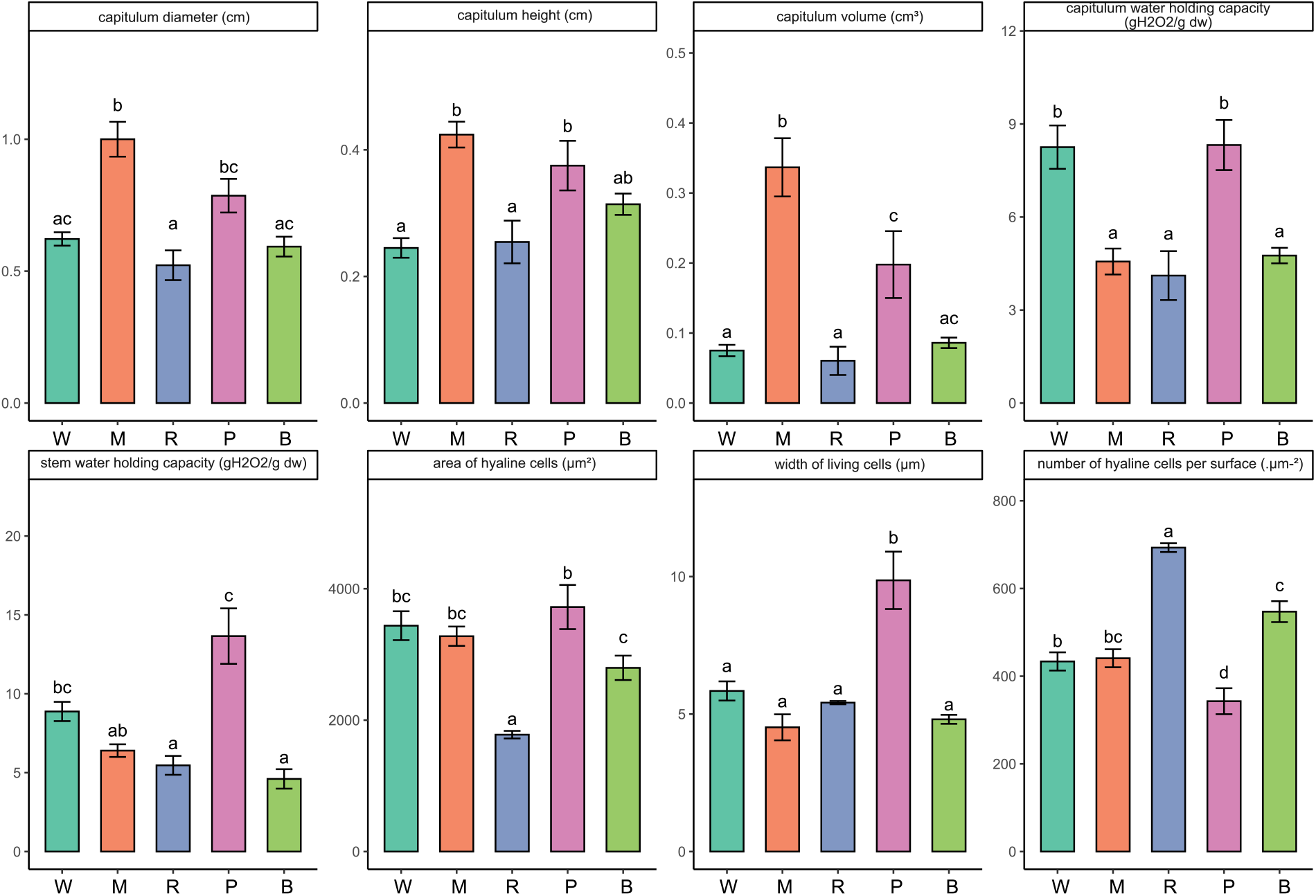
Barplot of anatomical traits of dominant *Sphagnum* mosses in five sites along the gradient. Data represents means and standard errors (SE). Letters indicate significant differences among warming treatments at p < 0.05 (linear mixed effect models). *Sphagnum* species are indicated by letter W = *S. warnstorfii*, M = *S. magellanicum*, R = *S. rubellum*, P = *S. papillosum*, B = *S. balticum.*

For *Sphagnum* enzymatic antioxidants and proteins, three groups were defined along the first NMDS axis: one group with S*. warnstorfii, S. magelanicum, S. balticum*, a second one with *S. rubellum* and a third one with *S. papillosum* (Figure 4c). These differences are driven by significant differences proteins and some enzymes (Figure 6). Proteins concentration (Figure 6; *F*_*4,16*_=5.33, *P*=0.0063, ANOVA) was the highest value in *S. magelanicum* and the lowest in *S. papillosum* (0.17 mg/ml and 0.072 mg/ml respectively). *S. papillosum* showed the highest APX activity with 27.94 mmol min^−1^ mg^−1^, while mean activity for the four other species was similar and around 12.1+− 2.37 mmol min^−1^ mg^−1^ (Figure 6; *F*_*4,16*_ = 6.9, *P* = 0.002, ANOVA). CAT and SOD showed similar trends with maximum activities in *S. papillosum* (Figure 6; CAT: *F*_*4,16*_ = 5.9, *P* = 0.0039; SOD: *F*_*4,16*_ = 9.52, *P* < 0.001, ANOVA). The POX activity was the highest in *S. rubellum* with 11.26 mmol min^−1^ mg^−1^, following by *S. papillosum* with 6.87 mmol min^−1^ mg^−1^, while for the rest almost no activity observed (Figure 6; *F*_*4,16*_=14.9, *P*<0.001, ANOVA).

**Figure 4.**
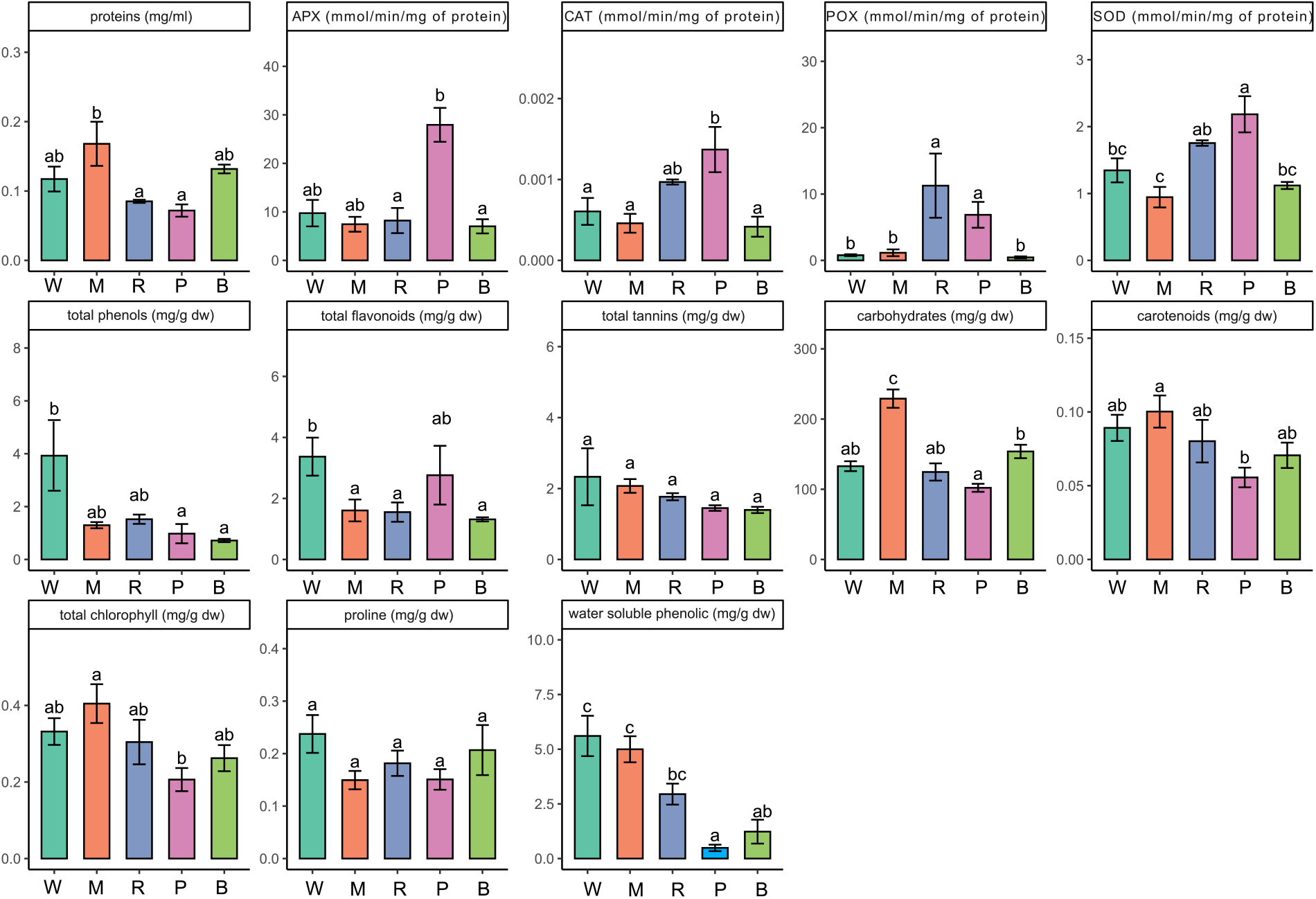
Barplot of *Sphagnum* enzymatic activities (a-e), metabolites and pigments (f-m) of dominant *Sphagnum* mosses in five sites along the gradient. Data represents means and standard errors (SE). Letters indicate significant differences among the sites along a latitudinal gradient at p < .05 (linear mixed effect models). *Sphagnum* species are indicated by letter W = *S. warnstorfii*, M = *S. magellanicum*, R = *S. rubellum*, P = *S. papillosum*, B = *S. balticum.*

For *Sphagnum* metabolites and pigments, three groups were identified along the second NMDS axis (Figure 4d): one group with S*. warnstorfii,* a second with *S. rubellum, S. papillosum, S. balticum* and a third group with *S. magelanicum*. Again, these patterns are explained by significant differences at the individual compound level. Carbohydrates had a maximum concentration in S*. magelanicum* (229.08 mg/g DW), and a minimum in *S. papillosum* (102.07 mg/g; Figure 6; *F*_4,16_=18.6, *P*<0.01, ANOVA). Chlorophyll (a+b) and carotenoids concentrations followed similar trends with the highest values in *S. magelanicum* and the lowest in *S. papillosum* (Figure 6; chl: *F*_4,16_=3.1, *P*=0.04; car: *F*_4,16_=2.9, *P*=0.052, ANOVAs). Phenolic concentrations followed somehow a latitudinal trend, with the highest value in *S. warnstorfii* (5.61 mg/g DW) and the lowest in *S. papillosum* (0.49 mg/g DW; Figure 6; *F*_4,16_ = 18.9, *P* < 0.001, ANOVA). Total polyphenols had the highest value in *S. warnstorfii*, while the lowest was observed in *S. balticum* (Figure 6j; *F*_4,16_=5.26015, *P*=0.0067). Total flavonoids also differed among *Sphagnum* species (Figure 6; *F*_*4,16*_ = 3.1, *P* = 0.046, ANOVA), with the lowest values found in *S. balticum*, *S. magellanicum* and *S. rubellum* and the highest in *S. warnstorfii*. Finally, concentrations of tannins and proline did not reveal any significant variation in their concentrations (Figure 6; *P*>0.05, ANOVAs).

### Linkages between Sphagnum attributes and environmental characteristics

The MFA of the three *Sphagnum* trait data sets and environmental and vegetation data sets confirmed the existence of an overall division among *Sphagnum* species attributes driven by environmental conditions (Table 1). Both *Sphagnum* anatomical traits and metabolites were significantly linked to *Sphagnum* proteins and enzymatic activities, environmental conditions and vegetation composition (Table 1). In addition, *Sphagnum* enzymatic activities and proteins were linked to vegetation composition (Table 1).

**Table 1.** RV-coefficients (below diagonal) and corresponding P values (above diagonal) among the *Sphagnum* attributes and environmental data used in the MFA of the five dominant species.

|  | anatomical | metabolites | enzymes | vascular | env-clim |
| --- | --- | --- | --- | --- | --- |
| anatomical |  | 0.085 | 0.006 | 0.005 | 0.002 |
| metabolites | 0.21 |  | 0.024 | 0.0004 | 0.0002 |
| enzymes | **0.29** | **0.260** |  | 0.014 | 0.57 |
| vascular | **0.30** | **0.390** | **0.260** |  | 0.0001 |
| env-clim | **0.33** | **0.410** | 0.090 | **0.50** |  |

In the RDA, the five *Sphagnum* species were clearly separated and revealed common responses of *Sphagnum* attributes to environmental variables (Figure 7). The model explained 41% (adjusted *R*^*2*^) of the variability in *Sphagnum* attributes. Flavonoids, water-holding capacities, enzymatic activities, width of chlorocystes were positively correlated to shrubs relative abundance and negatively to WTD and herbaceous relative abundance. Pigments, proteins, carbohydrates, and phenols were positively correlated to WTD, herbaceous relative abundance, temperature, annual precipitation, while negatively correlated to pH. *Sphagnum* anatomical traits such as capitulum volume, size and height were positively related to annual precipitations.

**Figure 5.**
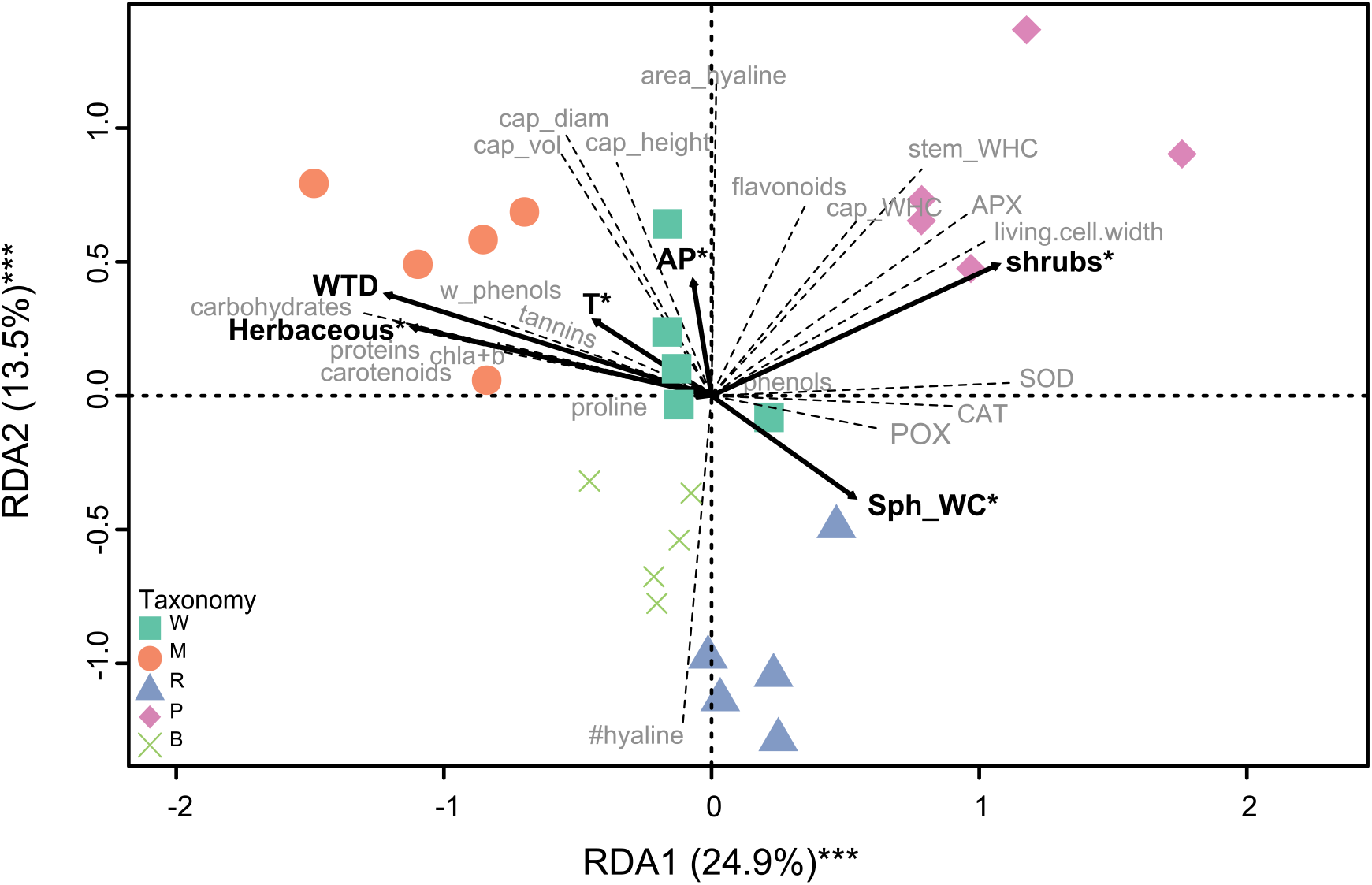
Redundancy analyses biplots (axes 1 and 2) of *Sphagnum* attributes from five dominant *Sphagnum* species. *Sphagnum* species are indicated by letter W = *S. warnstorfii*, M = *S. magellanicum*, R = *S. rubellum*, P = *S. papillosum*, B = *S. balticum*. Abbreviations: Sph_WC = S*phagnum* water content, AP = annual precipitation, T = the highest temperature of the warmest quarter, WTD = water table depth. Asterisks indicate significant *P*-values after permutation tests; *, *P* < 0.05, **, *P* < 0.01.

## Discussion

Our aim was to test whether phylogeny, taxonomy and environmental conditions were important determinants of *Sphagnum* anatomical and biochemical traits along a latitudinal gradient. Our screening of a broad range of *Sphagnum* anatomical and biochemical traits along a latitudinal gradient showed that several patterns emerged along with changing regional climate and local biotic/abiotic conditions. However, the majority of *Sphagnum* traits variation along the latitudinal gradient was inconsistent with the hypothesis that *Sphagnum* interspecific trait variability was driven by phylogeny. This indicates that regional climate and/or local biotic and abiotic conditions prevail on phylogeny in driving trait variations among *Sphagnum* mosses. Previous studies evidenced a strong phylogenetic conservatism among *Sphagnum* traits (Bengtsson et al., 2018, 2016; Limpens et al., 2017), especially among anatomical traits (Laine et al., 2020) However, most of these previous studies did not incorporated appropriate phylogenetic signal in their statistical analysis, as recently raised by Piatkowski and Shaw (2019), nor focused on the same traits and species. Our results are rather consistent with the recent findings from Piatkowski and Shaw (2019) who demonstrated that most *Sphagnum* traits related to litter quality and growth are not phylogenetically conserved. We show that *Sphagnum* metabolites and antioxidant enzymes are not phylogenetically conversed neither and strongly influenced by environmental conditions. This finding demonstrates that environmental filtering acts primarily not only on the taxonomic composition of *Sphagnum* mosses in peatlands (Robroek et al., 2017), but also on their biochemical machinery.

We found that the dynamic of *Sphagnum* traits were disconnected from the latitudinal gradient, and rather influenced by moisture conditions and vegetation community composition. This is in line with the contrasting effects of regional climate and/or local biotic and abiotic conditions on *Sphagnum* ecophysiology (Breeuwer et al., 2009; Gerdol, 1995; Gunnarsson et al., 2004; Robroek et al., 2007). We found that moisture conditions (WTD, annual precipitation and water content) were important determinants of *Sphagnum* anatomical traits, especially those related to water holding capacity. Such result was expected as *Sphagnum* species often have narrow habitat niches (Andrus et al., 1983; Johnson et al., 2015). *Sphagnum* species with different morphological and physiological constraints often replace each other along WTD gradient (Clymo and Hayward, 1982; Gong et al., 2019; Laing et al., 2014). The ability of *Sphagnum* to transport water to the capitula and retain cytoplasmic water are important traits controlling the moss community dynamic across peatland habitats (Bengtsson et al., 2020a; Gong et al., 2019; Hájek, 2020). Water-retention traits are key to supply sufficient moist in *Sphagnum* apical parts, minimize evapotranspiration (Rydin and McDonald, 1985; Titus and Wagner, 1984), and thus maintain important physiological processes such as photosynthesis (Bengtsson et al., 2020a; Gong et al., 2019; Jassey and Signarbieux, 2019). We evidenced that species with the highest water holding capacity also exhibited the highest antioxidant capacity, as showed by the high flavonoid and enzymatic activities in their tissues (Nakabayashi et al., 2014). As *Sphagnum* mosses lack stomata and water-conducting tissues, the ability to maintain moist in their active apical parts (capitula) rely on the water transporting efficiency from water table to the capitula and the desiccation avoidance (Clymo and Hayward, 1982). The desiccation avoidance strategy of *Sphagnum* mosses is mediated by large and empty hyaline cells but also requires strong cell walls to avert the collapse of large capillarity spaces (Hájek and Vicherová, 2014). The production of antioxidants, as protection of cell components from oxidative cell damages, is an important feature of desiccation tolerance in plants, and particularly in drought-hardening (Proctor, 1990). Synchronized action of major antioxidant enzymes (APX, CAT, SOD, POX) and specialized metabolites such as flavonoids outcome in detoxification of reactive oxygen species and limit oxidative stress in plants (Choudhury et al., 2013; Das and Roychoudhury, 2014; Noctor et al., 2018). These findings demonstrate that the degree of desiccation tolerance of *Sphagnum* mosses is not only limited to its morphological traits (Bengtsson et al., 2020a) but also rely on its proteome and metabolome to contain cell-damages.

On the contrary to what we expected, we did not evidence a strong effect of temperature on neither anatomical nor biochemical traits. Although our results showed higher polyphenol concentrations in *Sphagnum* tissues in warmer sites (Counozouls) compared to colder site (Abisko), such decrease in polyphenol content along with colder climate was most probably related to stronger nutrient limitation on plant growth in northern regions (Bragazza and Freeman, 2007; Dorrepaal et al., 2005) rather than a warming effect (Jassey et al., 2011a). Instead of a strong temperature effect, our study highlighted the importance of vascular plant functional types in driving *Sphagnum* biochemical traits, as showed by the increase in total tannins, total phenols, water soluble polyphenols, carbohydrates, pigments and proteins along with herbaceous cover. This suggests that *Sphagnum* mosses modulate their whole biochemical machinery in response to increasing competition with vascular plants. This agrees with previous findings showing that *Sphagnum* mosses shift their polyphenolic metabolism when resource competition with vascular plants for nutrients and light increases (Chiapusio et al., 2018). Indeed, *Sphagnum* mosses can compete with vascular plants through the release of specialized metabolites such as polyphenols that inhibit vascular plant germination and radicle growth (Chiapusio et al., 2013). The increase in total tannins and total phenols further indicates that *Sphagnum* produce a more decay-resistant litter. Such decay-resistant litter dampens microbial activity (Bengtsson et al., 2016) and increases peat accumulation rate. A lower microbial activity also induces low mineralization, which withdraws nutrients from the rhizosphere, hence reducing the capacity of vascular plants (Malmer et al., 2003). In parallel, the increase in carbohydrates, proteins and photosynthetic pigments suggest that *Sphagnum* produce structural and non-structural compounds to maintain its growth and enhance its stress tolerance to cope with vascular plant competition (Liu et al., 2019). In short, we suggest that vascular plant encroachment over *Sphagnum* causes alterations to the moss metabolome that directly impact *Sphagnum* functioning and fitness. While effects of vascular plants on *Sphagnum* fitness are detectable in the metabolome, they are missed entirely by morphological and anatomical traits. As such, considering the *Sphagnum* metabolome in understanding the response of *Sphagnum* mosses to environmental changes improves our mechanistic understanding of *Sphagnum* traits by providing intuitive explanations for *Sphagnum* performance where classical morphological and anatomical are lacking.

Our findings show that *Sphagnum* exposed to different local condition (WTD, vegetation composition) and global conditions (temperature, precipitation) initiate a diverse set of anatomical and biochemical responses, including developing traits for efficient water holding capacity, antioxidant activity and stress tolerance machinery, in order to maintain or survive changing conditions. These findings indicate that *Sphagnum* proteome, metabolome, and hence phenotype, underpin *Sphagnum* niche differentiation through their role in specialization towards biotic stressors, such as plant competitors, and abiotic stressors, such as temperature and drought, which are important factors governing *Sphagnum* growth (Weston et al., 2015). We suggest that these relationships represent previsouly unreported trade-offs of resource allocation by mosses, whereby *Sphagnum* mosses devote resources to maintain their growth to cope with environmental changes, or resist to increasing competition. We caveat that we measured only one species at a single site, and that our observations were undertaken at a single date. As such, future studies are needed to test the generality of the trade-offs detected here and their importance for processes of carbon cycling. Nevertheless, this study provided evidence that measurements of the proteome and metabolome, once properly incorporated with classical morphological and anatomical traits, may improve our understanding of the coupling between physiology and fitness in *Sphagnum* trait-based studies. Our findings illustrate the need of including biochemical traits in studies to better understand the response of *Sphagnum* species to environemntal changes, and reveal a further mechanism by which plant biochemicals assist stress tolerance adaptation and continious growth.

## Acknowledgments

This work was supported by the MIXOPEAT project (Grant No. ANR-17-CE01-0007 to VJ) funded by the French National Research Agency. We thank the *Plateforme Analyses Physico-Chimiques* from the Laboratoire Ecologie Fonctionnelle et Environnement (Toulouse) for their analyses (water extractable organic matter). BJMR was supported by the British Ecological Society research grant (SR17\1427).

